# Design, evaluation and implementation of synthetic isopentyldiol pathways in *Escherichia coli*

**DOI:** 10.1101/2023.02.21.529368

**Authors:** Yongfei Liu, Lin Chen, Pi Liu, Qianqian Yuan, Chengwei Ma, Wei Wang, Chijian Zhang, Hongwu Ma, An-Ping Zeng

**Affiliations:** Hamburg University of Technology, Institute of Bioprocess and Biosystems Engineering, Denickestr. 15, Hamburg 21073, Germany; Tianjin Institute of Industrial Biotechnology, Chinese Academy of Sciences, Tianjin 300308, China; Hua An Tang Biotech Group Co., Ltd, Guangzhou, China; Center of Synthetic Biology and Integrated Bioengineering, School of Engineering, Westlake University, Hangzhou, Zhejiang, 310024, China, 310024

**Keywords:** isopentyldiol, diol, retrosynthesis, pathway evaluation, thermodynamics analysis, synthetic biology

## Abstract

Isopentyldiol (IPDO) is an important raw material in cosmetic industry. So far IPDO is exclusively produced through chemical synthesis. Growing interest in natural personal care products has inspired the quest to develop a bio-based process. We previously reported a biosynthetic route that produces IPDO via extending leucine catabolism (route A), the efficiency of which, however, is not satisfactory. To address this issue, we computational designed a novel non-natural IPDO synthesis pathway (Route B) using RetroPath RL, the state of art tool for bio-retrosynthesis based on Artificial Intelligence methods. We compared this new pathway with route A and another two intuitively designed routes for IPDO biosynthesis from various aspects. Route B, which exhibits the highest thermodynamic driving force, least non-native reaction steps and lowest energy requirements appeared to hold the greatest potential for IPDO production. All three newly designed routes were then implemented in *E. coli* BL21(DE3) strain. Results show that the computationally designed route B can produce 2.2 mg/L IPDO from glucose, whereas no IPDO production from routes C and D. These results highlight the importance and usefulness of *in silico* design and comprehensive evaluation of the potential efficiencies of candidate pathways in constructing novel non-natural pathways for the production of biochemicals.

## Introduction

Presently, the majority of value-added compounds, such as solvents, biofuels and polymer precursors, are produced through petroleum-dependent and energy-intensive chemical processes. Great concerns about climate change, air pollution and oil price fluctuation have inspired the quest for alternative green processes that produce desired products with low toxic by-product formation and carbon dioxide emission. Bio-based production processes using renewable feedstocks as substrates hold a great potential for a sustainable **s**upply of chemicals. Synthetic biology aims at designing and altering the host metabolic system, as well as introducing artificial pathways whose behavior can be predicted, programed, and finally, controlled (Glass 2018; Lee and Kim 2015; Mitsuhashi 2014).

Short chain (C3-C5) diols are considered a class of important green platform chemicals that have wide applications in manufacturing a variety of commercial chemical products, such as cosmetics, feed additives, polymers, solvents and pharmaceuticals (Wang J et al. 2020; Zeng and Sabra 2011; Zhang et al. 2017). Isopentendiol (IPDO) is an appealing ingredient in cosmetic industry. It has no odor or irritation, and is evaluated as safe & eco-friendly material. Compared to other diols used in cosmetics (such as propylene glycol, butylene glycol or dipropylene glycol), IPDO presents better skin feel, and deodorant and antibacterial performances, therefore can works as moisturizer, solubilizer and booster of preservative. Recently, we proposed a general route for microbial production of structurally diverse diols by expanding amino acid metabolism (Liu Y. et al. 2022). This general route combines oxidative and reductive formations of the diol hydroxyl groups, and comprises four reactions catalyzed by amino acid hydroxylase, L-amino acid deaminase, α-keto acid decarboxylase and aldehyde reductase consecutively. As a proof of concept, the feasibility of IPDO production via this general route has been verified (named here as route A). To increase IPDO titer further efforts have been made, including screening for pathway enzymes with higher activity, enhancing the supply of the precursor amino acid, and directed evolution of the key enzyme hydroxylase. The achieved IPDO titer of 310 mg/L is, however, still far from the requirement of a practical application. In view of this, we explore in the present study the possibility of producing IPDO via alternative artificial biosynthetic routes.

Designing an artificial biosynthetic route is a process which generally consists of three steps: (1) design and evaluation of pathways; (2) verification of the designed pathways *in vivo* or *in vitro*; and (3) improving the production of target compounds by applying metabolic engineering strategies (Lee and Kim 2015). The development of computational pathway prediction tools has largely facilitated the identification of possible metabolic pathways and the corresponding enzymes. Currently, a number of prediction algorithms have been developed to aid in the biosynthetic route design, such as BINCE (Henry et al. 2010), RetroPath RL (Carbonell et al. 2014), and Retrobiocat (William Finnigan, 2021). These pathway design tools use the retrosynthetic analysis (RTSA) method often used in chemical synthesis. RTSA uses a set of reaction rules to backtrack from target products to potential reactants and connect individual reactions to form pathways. A scoring function is then used to rank the generated pathways based on reaction thermodynamic parameters, substrate/product similarity, potential toxicity of intermediates, and the number of enzymes that need to be heterologously expressed so as to select for more rational biosynthetic pathways. The advantage of RTSA based pathway design is that the target product is not limited to known natural metabolites of a microbial host but can also be a completely new molecular structure. Moreover, as the technology of enzyme design advances, the diversity of substrates that enzymes can catalyze will broaden, enabling the synthesis of more diverse products using the generalized reaction rules of the retrosynthetic pathways design method. One landmark in the application of computer-aided design in metabolic engineering is the biological production of 1,4-butanediol (1,4-BDO) (Yim et al. 2011). In their work, more than 10,000 pathways were initially identified using the software Bio-Pathway Predictor, followed by selection using a pathway selection software to sort and rank the pathways based on attributes including maximum theoretical yield, thermodynamic feasibility, number of non-native reaction steps, and pathway length. As the result, two high-ranking artificial routes were selected for metabolic engineering, and the best recombinant strain produced 18 g/L 1,4-PDO from glucose after 5 days of cultivation.

Prior to experimental implementation of the three designed routes, we investigated the potential of these designs by using two *in silico* analysis tools. For comparison, route A is served as the reference route and evaluated with the other three routes in parallel. We used the flux balance analysis (FBA) (Orth et al. 2010) to predict the maximum IPDO yield and the max-min driving force (MDF) (Noor et al. 2015) to determine the pathway thermodynamics so as to predict the feasibility and to identify the key bottlenecks in these pathways. Both tools are well-elucidated and can be easily implemented without requirement of experimental data, thereby providing a simple and effective means of evaluating different artificial pathways that produce the same target compound. Afterwards, in comparison with route A all three artificial biosynthetic routes were separately introduced into *E. coli* BL21(DE3) strain, and their performances on IPDO production were analyzed and compared with the *in silico* prediction results.

## Materials and methods

### 1. Strains, plasmids, chemicals and cultivation conditions

Strains and plasmids used in this work are listed in Table S1. *E. coli* Top10 was used for gene cloning and *E. coli* BL21(DE3) for protein expression. Primers used in this work are listed in Table S2. Genes encoding α-ketoisocaproate dioxygenase, i.e. SnHPD from of *Streptomyces noursei* (DSM 40635) and AaHPD from *Allokutzneria albata* (DSM 44149), and methylglyoxal reductase GRE2 from *Saccharomyces cerevisiae* (strain S288c) were amplified from the corresponding genome DNA. Genes encoding carboxylic acid reductase CAR from *Nocardia iowensis*, leucine hydroxylase MFL from *Methylobacillus flagellatus KT*, L-amino acid deaminase AAD from *Proteus vulgaris*, glycerol dehydratase GDHt from *Klebsiella pneumonia* and aldehyde-deformylating oxygenase ADO from *Cyanobacteria* were synthesized by the Genscript Company (Netherlands). Genes encoding α-keto acid decarboxylase KDC from *Lactobacillus lactis* was previously synthesized in our lab (Zhang et al. 2019). <u>Mevalonate and IPDO were purchased from the Sigma-Aldrich Company (Germany).</u> 3-hydroxyisovalerate (3-OH-IV), α,β-dihydroxyisovalerate (α,β-DHIV), α-ketoisocaproate (α-KIC) were purchased from the TCI Company (Japan). Isopropyl β-D-1-thiogalactopyranoside (IPTG) was purchased from the Carl Roth Company (Germany). 4-hydroxyleucine (4-OH-Leu) was purchased from the AKos GmbH (Germany).

For shake flask culture, individual colonies were picked up from agar plates and cultivated in the LB medium (5 g/L yeast extract, 10 g/L tryptone, and 10 g/L NaCl) overnight. Harvested cells were then used to inoculate the fermentation medium II (FM-II medium containing 30 g/L glucose, 0.5 g/L MgSO_4_·7H_2_O, 3 g/L KH_2_PO_4_, 12 g/L K_2_HPO_4_, 4 g/L (NH_4_)_2_SO_4_, 1 g/L yeast extract, 2 g/L monosodium citrate and 0.1 g/L FeSO_4_·7H_2_O, pH 7.0) at the inoculation rate of 1: 100 (v/v). After OD_600_ reached 0.4 ∼ 0.6, 0.1 mM IPTG was added to induce protein expression, and the cultivation was carried out at 30°C, 200 rpm for 48 h. Samples were collected by centrifugation and the supernatants were filtered with a filter membrane (0.22 μm) and stored at -20°C for subsequent analysis by gas chromatography (GC) or gas chromatography-mass spectrometry (GC-MS).

### 2. Plasmid construction

Taking the pET-mfl as an example, the procedure of plasmid construction was as follows: The linearized *mfl* fragment with 15 bp extension homologous at its ends was obtained by using primers pET-mfl-LF and pET-mfl-LR with 50 ng of synthesized *mfl* as template. The pET28a vector was linearized by using primers pET-vec-LF and pET-vec-LR with 50 ng of pET28a plasmid as template. The linearized pET28a vector and *mfl* fragment were ligated using the In-Fusion HD Cloning Kit (Clontech, Takara, Japan), and the ligated product was then transformed into *E. coli* Top10 strain and cultivated on LB agar plate containing 50 mg/L kanamycin (Kan) at 37°C overnight. The next day, colony PCR was applied to verify positive colonies, and the plasmids were extracted to sequence for further verification.

### 3. Computational pathway design

In order to design possible IPDO pathways, we performed a retrosynthesis analysis of IPDO using the software Retropath RL. First, we made each intermediate of the designed routes have very high structural similarity to the structure of a corresponding natural intermediate metabolite. The substrate similarity cutoff value was set to 0.5, width to 10, depth to 7, max rollout to 10, reaction rule radius to 2, 4, 8, and 16, score function to biological pathway scoring. We picked the *E. coli* intermediate metabolite database available in their software as Sinks so that retrosynthesis search will be stopped when a metabolite synthesizable in *E. coli* is reached. Then, to find more potential synthetic pathways, we varied the calculation parameters, reduced the substrate similarity to 0.1, switched to a random scoring method, and expanded the source of metabolites to all detectable intermediate metabolites.

### 4. Flux balance analysis of IPDO biosynthesis via four different artificial routes

The optimal metabolic flux distributions of the strains carrying different IPDO synthetic pathways were calculated using flux balance analysis (FBA) and the genome-scale metabolic model iY75_1357 for *Escherichia coli* str. K-12 substr. W3110 (http://bigg.ucsd.edu/models/iY75_1357). The reactions in each artificial route were as follow:

#### Route A

1. Leu + α-KG + O_2_ → 4-OH-Leu + succinate + CO_2_
2. 4-OH-Leu + 1/2 O_2_ → 4-OH-α-KIC + NH_3_
3. 4-OH-α-KIC → 3-OH-3-M-BA + CO_2_
4. 3-OH-3-M-BA + NADPH + H^+^ → IPDO + NADP^+^

#### Route B

1. α-KIC + O_2_ → 3-OH-IV + CO_2_
2. 3-OH-IV + ATP + NADPH + H^+^ → 3-OH-3-M-BA + AMP + Ppi + NADP^+^
3. 3-OH-3-M-BA + NADPH + H^+^ → IPDO + NADP^+^

#### Route C

1. α, β-DHIV + ATP + NADPH + H^+^ → 2,3-diOH-IVA + AMP + Ppi + NADP^+^
2. 2,3-diOH-IVA + NADPH + H^+^ → 3-M-BTO + NADP^+^
3. 3-M-BTO →3-OH-3-M-BA + H_2_O
4. 3-OH-3-M-BA + NADPH + H^+^ → IPDO + NADP^+^

#### Route D

1. acetoacetyl-CoA + acetyl-CoA + H_2_O → HMG-CoA + CoA
2. HMG-CoA + 2NADPH + 2H^+^ → MVA + CoA + 2NADP^+^
3. MVA + ATP + NADPH + H^+^ →3,5-diOH-3-M-PA + AMP + Ppi + NADP^+^
4. 3,5-diOH-3-M-PA + 2NADPH + O_2_ + 2H^+^→ IPDO + HCOOH + H_2_O + 2NADP^+^

The resulting metabolic sub-models were named as iY75_1357_IPDO_A, iY75_1357_IPDO_B, iY75_1357_IPDO_C and iY75_1357_IPDO_D and used for a constraint-based FBA, in which the glucose uptake rate was set at 15 mmol/g/DCW and the source of oxygen, nitrogen, phosphate and sulfur were not constrained.

### 5. Thermodynamic analysis of the four IPDO synthetic routes

For analyzing the thermodynamic efficiency of the four IPDO synthetic routes, Gibbs free energies of reactions (Δ_r_G’^0^) were calculated using the component contribution method (Jankowski et al. 2008). The max-min driving force (MDF) analysis was performed using the Python packages equilibrator-api and equilibrator-pathway (Flamholz et al. 2012). The concentration boundaries of metabolites were set with reference to the literature reports (Yang et al. 2021) as follows: ATP : 1 mM; ADP: 0.1 mM; Pi: 0.1 mM; NADPH: 1 mM; NADP: 0.1 mM; AcCoA: 10 mM; AaCoA: 0.0005-10 mM; CoA: 0.0005-10 mM; IPDO: 0.0005-10 mM; 3HmCoA: 0.0005-10 mM; HS_CoA: 0.0005-10 mM; MVA: 0.0005-10 mM; 35dmp: 0.0005-10 mM; formate: 0.0005-10 mM.

### 6. Enzyme expression and purification

For the expression and purification of each target protein, single colony from LB agar plate was inoculated into 5 mL LB medium at 37°C, 220 rpm overnight. The overnight culture was then inoculated into 50 mL LB medium in 300-mL shake flasks. After growing at 37°C, 220 rpm to an OD_600_ of 0.4 ∼ 0.6, 0.1 mM IPTG was added into the medium to induce the enzyme expression, and the cultivation was continued at 30°C, 220 rpm for 12 ∼16 h. After cooling down on ice for 30 min, cells were harvested by centrifugation at 4°C, 5000 rpm and washed with 30 mL of a buffer containing20 mM sodium phosphate, 500 mM NaCl and 20 mM imidazole (pH 7.4). Cell pellets were re-suspended in 3 mL of the same buffer and disrupted by the multidirectional, simultaneous beating with specialized lysing matrix beads in a FastPrep® 24 homogenizer (MP Biomedicals). The supernatant containing the target protein was collected by centrifuging at 4°C, 13,000 rpm for 20 min. The target protein was purified by loading the supernatant onto a prepacked His SpinTrap column (GE Healthcare) and eluting with a buffer containing 20 mM sodium phosphate, 500 mM NaCl and 500 mM imidazole (pH 7.4), followed by buffer change into a HEPES buffer (pH 7.5) using the Amicon® Ultra-0.5 centrifugal filter device (10K MWCO) at 4°C for the subsequent enzyme assay. The concentration of the purified protein was determined according to Bradford’s method using bovine serum albumin (BSA) as standard.

### 7. Enzymatic activity assays

Assay of the carboxylic acid reductase (CAR) activity was performed as described previously (Matthew Moura 2015) with minor modification. The enzyme activity was measured by monitoring the disappearance of NADPH via absorbance at 340 nm. The reaction mixture was composed of 80 mM HEPES buffer (pH 7.5), 10 mM MgCl_2_, 0.25 mM NADPH, 1 mM ATP, 5 mM of a carboxylic acid substrate and 100 μg purified CAR in a total volume of 0.2 mL. The reaction mixture without the substrate was incubated to 30°C, and the reaction was started by adding the corresponding carboxylic acid and allowed to carry out for 5 to 15 min. The enzymatic activity was determined by monitoring the disappearance of NADPH by measuring the absorbance at 340 nm.

### 8. Whole-cell catalyzed conversion of α-KIC to 3-OH-IV

For HPD expression, corresponding genes of two α-ketoisocaproate dioxygenase (HPD), namely SnHPD from *Streptomyces noursei* and AaHPD from *Allokutzneria albata*, were separately inserted into the pET22b vector and transformed into *E. coli* BL21(DE3) to yield recombinant strains IPB03 and IPB04, respectively. IPB03 and IPB04 were cultivated in LB medium containing 0.1 mM IPTG to induce the expression of the HPDs. After cultivated overnight, cells were harvested by centrifugation at 4°C, 5000 rpm for 10 min, washed with 0.9% NaCl, and then re-suspended in 5 mL of 0.1 M Tris-buffer (pH 6.5) containing 33 mM α-KIC, 3 mM FeSO_4_, 1 mM DTT, and 5 mM ascorbate. The whole-cell catalysis of α-KIC conversion to 3-OH-IV was carried out at 30°C and 220 rpm in 100-mL baffled shake flask. Samples were taken after 6 h and 24 h of incubation to measure the concentration of remaining 3-OH-IV in the supernatant.

### 9. Analysis of IPDO by GC and GC-MS

To measure the amount of IPDO in fermentation broth, 50 μL of filtered culture supernatant was mixed with 100 μL of acetonitrile (ACN) and 50 μL of a derivatization reagent (comprised of 5 mg/mL phenylboronic acid dissolved in 2,2-dimethoxypropane) in a 1.5 mL tube, and the derivatization was performed by shaking at 600 rpm at room temperature for 15 min. After centrifugation to remove the precipitated protein the supernatant was transferred into a glass vial for subsequent GC analysis.

GC analysis of the derivatized IPDO was achieved on a phenomenex HP-5 capillary column (30 m × 0.25 mm × 0.25 µm) using a Varian 3900 GC instrument equipped with a flame ionization detector (FID). 1 µL of the derivatized sample was injected at 270°C using a splitless/split injector (splitless for 1 min, then split at a ratio of 1:10). Nitrogen gas was used as the carrier gas at a flow rate of 1.5 mL/min. The column oven temperature program was as follows: hold at 100°C for 2 min, then ramped at 10°C/min to 220°C, and finally ramped at 20°C/min to 260°C and held for 5 min. The FID detector was operated at 300°C.

GC/MS analysis of the derivatized IPDO was carried out using an Agilent 7890B gas chromatograph equipped with an Agilent DB-5ms UI non-polar capillary column (30 m × 250 μm × 0.25 μm) and coupled to an Agilent MSD 5977 mass spectrometer. 0.2 µL of sample was injected into a Gerstel CIS injector at a split ratio of 1:5. The initial operating temperature was 60°C (hold for 0.1 min), then ramped at a rate of 12°C/s to 270°C and held for 1 min. The column oven was held at an initial temperature of 60°C for 2 min, then ramped to 270°C at 20°C/min and held for 12 min. The MS was performed in scan mode in the range of 2 ∼ 500 amu, with a solvent delay of 4.5 min. The confirmation of IPDO was performed through the comparison of the mass spectra acquired as well as the retention times between the sample and the IPDO standard.

## Results

### 1. Design and evaluation of the novel IPDO biosynthetic routes

To find a potential retrosynthetic pathway, we used the bio-retrosynthesis analysis software RetropathRL to make a computational prediction of the biosynthetic pathway for the target product IPDO. The prediction gave only one reasonable biosynthetic pathway: Firstly, α-KIC was oxidatively decarboxylated by 4-hydroxyphenylpyruvate dioxygenase/α-ketoisocaproate dioxygenase (HPD) (EC: 1.13.11.27) to produce 3-OH-IV (Figure 1). 3-OH-IV is then converted to 3-hydroxy-3-methylbutanal (3-OH-3-M-BA) by a carboxylic acid reductase (CAR) (EC: 1.2.1.30). Subsequently, 3-OH-3-M-BA is reduced to IPDO by 1,3-propanediol dehydrogenase (PDDH) (EC: 1.1.1.202). Another pathway was obtained from the second round of selection after modification of the parameters (Figure S1). In this route, 3-OH-IV is obtained by hydrolysis of ethyl 3-hydroxy-isovalerate (E-3-OH-IV), which is picked as the starting metabolite of the designed route. However, considering that this precursor is not a natural metabolic intermediate of *E. coli* but a structure deduced by retrosynthesis, we chose the former pathway for further research.

**Figure 1.**
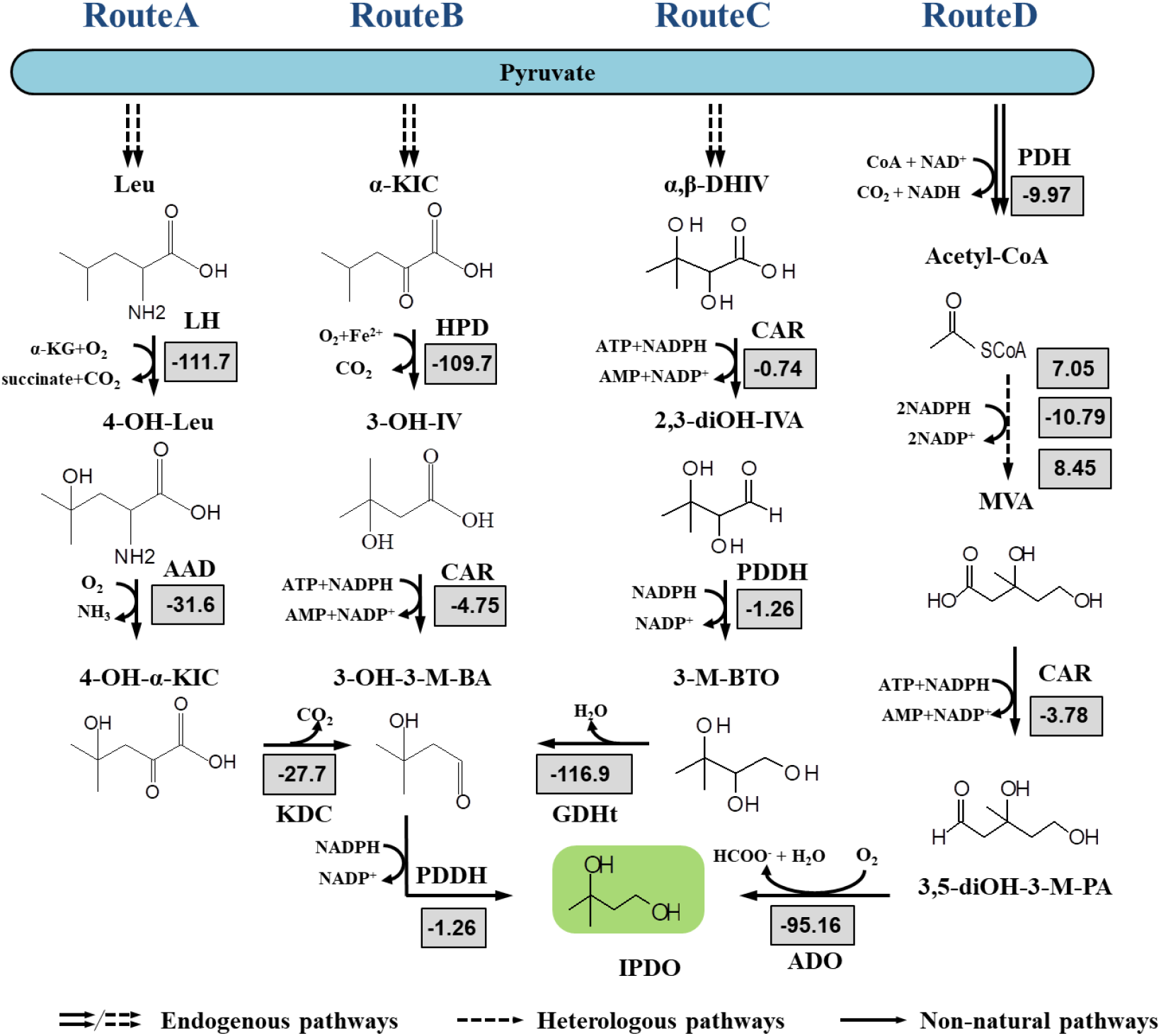
Proposed synthetic routes for IPDO production. All four routes produce IPDO by extending endogenous intermediate metabolisms of *E. coli*. Route A and route B divert flux from the leucine synthesis pathway and are composed of four and three exogenous reaction steps, respectively. Route C starts from α, β-dihydroxyisovalerate and consists of four non-native reaction steps. In route D, IPDO is synthesized by extending the mevalonate pathway via two exogenous reaction steps. The standard Gibbs free formation energy of every non-natural product was estimated using the group contribution method (Jankowski et al. 2008). Abbreviations: LH, leucine hydroxylase; AAD, L-amino acid deaminase; KDC, α-keto decarboxylase; PDDH, 1,3-propanediol dehydrogenase; HPD, α-ketoisocaproate dioxygenase; CAR, carboxylic acid reductase; GDHt, glycerol dehydratase; ADO, aldehyde-deformylating oxygenase; 4-OH-Leu, 4-hydroxyleucine; 4-OH-α-KIC, 4-hydroxy-α-ketoisocarproate; 3-OH-3-M-BA, 3-hydroxy-3-methylbutanal; α-KIC, α-ketoisocaproate; 3-OH-IV, 3-hydroxyisovalerate; α, β-DHIV, α, β-dihydroxyisovalerate; 2,3-diOH-IVA, 2,3-dihydroxy-3-methylbutanal; 3-M-BTO, 3-methyl-1,2,3-butanetriol; Pyr, pyruvate; MVA, mevalonate; 3,5-diOH-3-M-PA, 3,5-dihydroxy-3-methylpentanal; IPDO, isopentyldiol.

We notice that one distinct structural feature of IPDO is that it contains an isobutyl group, which motivates us to develop novel biosynthesis pathways that generate IPDO from metabolites that share the similar structure. In this regard, we propose two new biosynthetic routes (route C and D) by selecting α, β-dihydroxyisovalerate (α, β-DHIV), an intermediate of valine biosynthesis pathway, and mevalonate (MVA), respectively, as the precursors for IPDO production. Specifically, in route C, CAR catalyzes the reduction of α, β-DHIV to generate 2,3-dihydroxy-isovaleraldehyde (2,3-diOH-IVA), which in turn is converted to 3-methylbutane-1,2,3-triol (3-M-BTO) by PDDH. 3-M-BTO is transformed to 3-OH-3-M-BA by glycerol dehydratase (GDHt, EC: 4.2.1.30) and subsequently to IPDO by PDDH. In route D, IPDO is produced by stretching the well-elucidated heterologous MVA catabolism (Wang et al. 2016b; Ye et al. 2017). Specifically, CAR is employed to reduce MVA to 3,5-dihydroxy-3-methylpentanal (3,5-diOH-3-M-PA), which is subsequently converted to IPDO by aldehyde-deformylating oxygenase (ADO) (EC: 4.1.99.5). Unfortunately, routes C and D cannot be predicted using the bio-retrosynthesis analysis software RetropathRL even after modification of the parameters. The reasons behind might be that the rule set used by retropathRL is too restrictive in the types of non-reactive atoms in the molecular it uses for similarly structured substrates, consequently restricting the application potential of its rules and leading to a limitation of the algorithm’s ability to design new non-natural pathways.

Then, to evaluate the potential efficiencies of the four routes we set several criteria including theoretical yield of IPDO, pathway thermodynamics estimated by the max-min driving force (MDF), energy efficiency in form of ATP and NADPH cost, number of non-native steps, and penalties posed by inappropriate properties of intermediates and pathway enzymes (Table 1). We used FBA to predict their theoretical IPDO yields and corresponding metabolic flux distributions. Results showed that all three proposed IPDO pathways have higher IPDO theoretical yields than our previously reported route A (Table 1). The low yield (0.628 mol/mol glucose) of route A is due to the fact that hydroxylation of leucine into 4-OH-Leu in route A is coupled with the oxidation of α-KG to succinate and reduction of O_2_ to CO_2_. This means that in addition to the carbon loss in form of CO_2_, a certain metabolic flux has to be shunted into α-KG formation to make this coupled reaction feasible, which may impose an “additional burden” for the efficient IPDO production via this route. In contrast, route C starting from α,β-diOH-IV without carbon loss gives the highest IPDO theoretical yield, reaching 0.827 mol/mol glucose (Table 1, Figure S2), which is 96.5% of the maximal theoretical yield (0.857 mol/mol glucose) determined purely by the energy balance (Dugar and Stephanopoulos 2011). The theoretical yields of IPDO of routes B reaches 0.667 mol/mol glucose. Specifically, one carbon atom is lost as carbon dioxide (CO_2_) in the oxidative decarboxylation of α-KIC to 3-OH-IV which reduces its IPDO theoretical yield. As to route D, there are two reactions that involve the carbon loss: one is the conversion of pyruvate to acetyl-CoA, accompany with CO_2_ formation, and the other one is the last reaction step of converting 3,5-diOH-3-M-PA to IPDO by ADO, giving rise to a theoretical IPDO yield of 0.667 mol/mol glucose.

**Table 1.** Criteria for evaluating the potential efficiencies of four IPDO synthetic routes

| Criteria |  | RouteA | RouteB | RouteC | RouteD |
| --- | --- | --- | --- | --- | --- |
| Theoretical yield (mol/mol glucose) |  | 0.628 | 0.667 | 0.827 | 0.667 |
| MDF values (kJ/mol) |  | 31.54<sup>a</sup> | 31.54<sup>a</sup> | 6.96<sup>a</sup> | 38.37<sup>a</sup> |
|  |  | 84.67<sup>b</sup> | 74.93<sup>b</sup> | --- | --- |
| NAD(P)H required |  | 1 | 2 | 3 | 3 |
| ATP required |  | 0 | 1 | 1 | 1 |
| Number of non-native steps |  | 4 | 3 | 4 | 5 |
| Penalty | Intermediate | | α-KIC<br>(low affinity to HPD) | $\alpha,\beta$-DHIV<br>(low affinity to CAR) | |
|  | Enzyme |  |  | GDHt<br>(cofactor: VB<sub>12</sub>) | ADO<br>(unclear mechanism) |
| Verification |  | Yes | Yes | No | No |

Note: <sup>a</sup>: MDF values of all reactions in IPDO synthesis routes.
<sup>b</sup>: MDF values of the rest reactions after removing the last common rate-limiting reaction catalyzed by PDDH in routes A and B.

MDF is used to quantify a pathway’s tendency to operate near-equilibrium and is calculated by setting the minimal driving forces of all reactions as the optimization goal and maximizing it (Noor et al. 2015). MDF determines the magnitude of the thermodynamic bottleneck of a given route. In principle, a route with a higher MDF value always means it has a less severe thermodynamic bottleneck and is thermodynamically more favorable. As also shown in Table 1, we determined the MDF values for all the four IPDO synthetic routes under the given conditions (Section 4 in the “material and method”) to evaluate their thermodynamic favorability. Route D with a MDF value of 38.37 kJ/mol seems to be thermodynamically most favorable (Table 1, Figure 2) among the four routes. However, it has to be noted that its higher MDF value is mainly contributed by the high Δ_r_G’ (−95.16 kJ/mol) of the last reaction catalyzed by ADO (Figure 1, Figure 2), which indicates that all upstream reactions in route D are rate-limiting reactions (Figure 2). This will practically be a problem for the subsequent optimization of this route. Route C gives the lowest MDF value (6.96 kJ/mol), which is mainly attributed to the low Δ_r_G’ values of the three reactions catalyzed by CAR (−0.74 kJ/mol), PDDH (−1.26 kJ/mol) and PDDH (−1.26 kJ/mol) consecutively. In comparison, the MDF values of routes A and B are equal and lower than that of route D. However, since routes A and B also share a thermodynamically clearly unfavorable reaction, i.e. the last reaction catalyzed by PDDH (Δ_r_G’: -1.26 kJ/mol, Figure 1, Figure 2), it seems that their MDF values are severely constrained by this bottleneck reaction. In this way, the contribution of other reactions to the thermodynamic driving forces of the routes is overwhelmingly obscured. To this end, we removed this common bottleneck reaction and recalculated the MDF values of the rest reactions in routes A and B, which resulted in higher MDF values for both routes (84.67 kJ/mol *vs* 74.93 kJ/mol). Moreover, the resulting cumulative Δ_r_G’ values in routes A and B demonstrate a nearly linear decrease with each added reaction. The thermodynamically highly favorable reactions of hydroxylation, deamination and decarboxylation in route A and oxidative decarboxylation reaction in route B provide the powerful push needed for the thermodynamically unfavorable reduction formation of hydroxyl group in the last step. This indicates that in pathway design, it is important to consider the thermodynamic topology of the reactions in an envisaged synthetic pathway.

**Figure 2.**
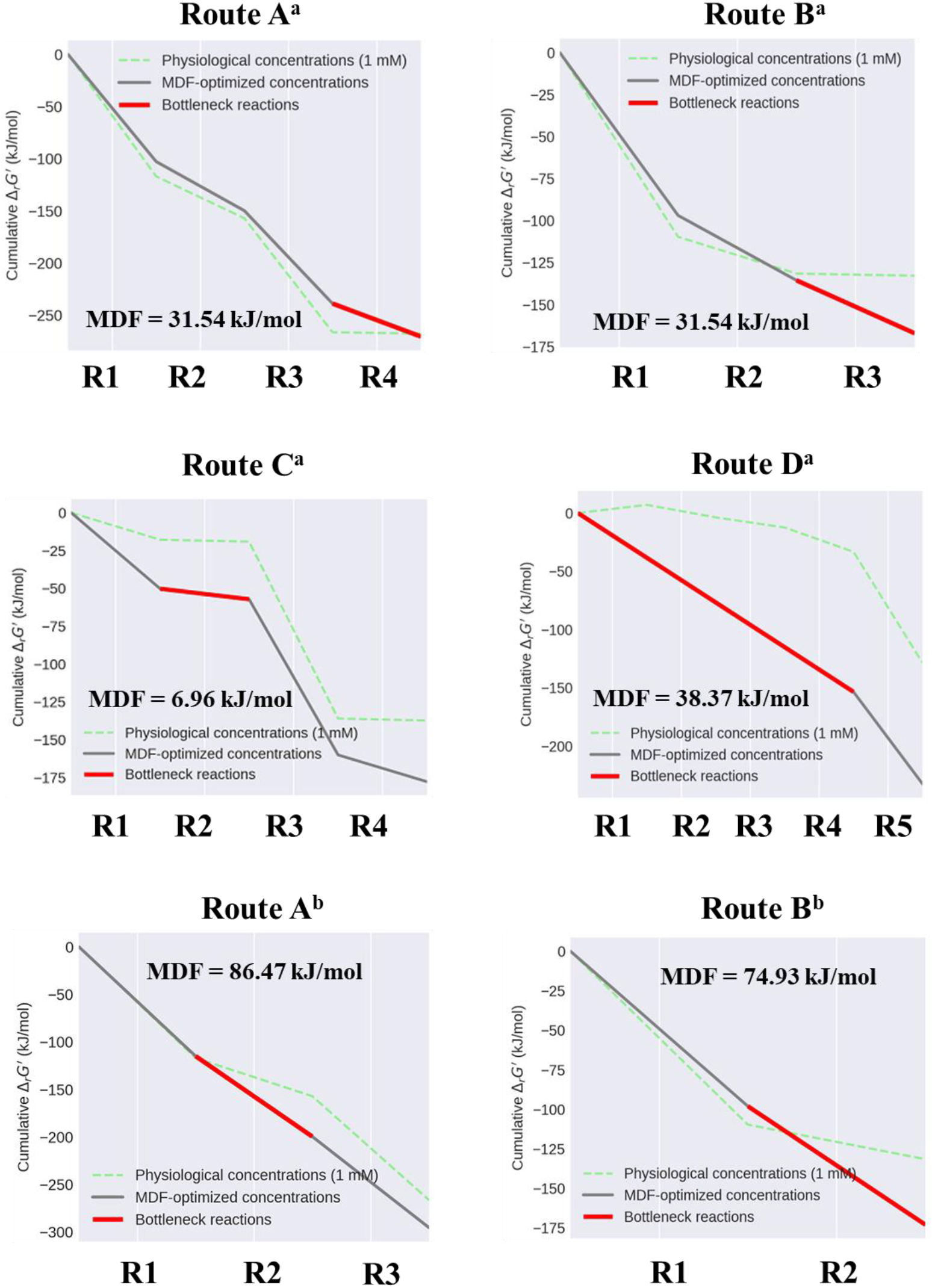
MDF values of four IPDO biosynthesis routes. Green dot lines represent the cumulative Δ_r_G’ values of reactions in IPDO biosynthesis routes when the concentrations of all substrates are set as 1 mM, while solid lines represent the cumulative Δ_r_G’ values of reactions when the concentrations of all substrates are under the optimal conditions evaluated by MDF. The reactions with the lowest Δ_r_G’ values are regarded as bottleneck reactions (as shown in red solid line in Figure 2). Note: ^a^: MDF values of all reactions in the IPDO synthesis routes. ^b^: MDF values of the rest reactions after removing the last common rate-limiting reaction in routes A and B.

The properties of intermediates and enzymes should also be taken into consideration to evaluate the potential of non-natural biosynthetic pathways. In many artificial biosynthetic pathways, some metabolites are produced as the outcome of promiscuous activity of heterologous enzymes. The physiochemical properties of these metabolites may restrict the structure of the pathway and ultimately constrain the product yield. Major physiochemical properties of intermediates to be considered include toxicity, stability and permeability (Bar-Even et al. 2012). In addition to high catalytic activity and specificity towards target substrate, an ideal enzyme with industrial application prospects would have clearly known catalytic mechanism, is easy and cheap to prepare and use, and is resistant to fluctuating cultivation environment (temperature or pH etc.). Accordingly, we checked the intermediates and enzymes involved in the four IPDO routes, and regarded the unfavorable intermediates and enzymes in these four routes as “penalties”. In the aspect, glycerol dehydratase GDHt employed in route C requires the expensive vitamin B_12_ as a cofactor to maintain its activity, which may largely limit its industrial application. The catalytic mechanism of aldehyde-deformylating oxygenase ADO in route D is still controversial (Aukema et al. 2013; Jia et al. 2015; Pandelia et al. 2013; Wang et al. 2016a), making its subsequent rational engineering for further improvement of its activity challenging. The drawbacks of GDHt and ADO reduce the implementation priority of route C and route D.

Overall, we came up with three new IPDO biosynthetic routes and compared their potential efficiencies with our previously reported route A. All three new routes show higher theoretical IPDO yields than route A, implying greater potential from a glucose utilization perspective. Route A and B, which produce IPDO by combing oxidative and reductive pathways of -OH group formation, require fewer reducing powers and exhibit higher thermodynamic driving force (as shown by their higher MDF values and bottleneck reactions determined by MDF) than route C and D that generate both hydroxyl groups of IPDO through reductive pathway. In addition, route B gives the fewest non-native reaction steps among all four IPDO routes. Unlike route C and D, route B doesn’t employ enzymes whose activities are condition-dependent or not well-characterized. All these advantages highlight the priority of route B.

In the follow-up study, we conducted the implementation of all three newly proposed IPDO biosynthetic routes in *E. coli*, so as to explore the effectiveness of the methods and criteria used in this study for a priori *in silico* estimation of the thermodynamic feasibility of designed pathways for a given product and the relative likelihood of alternative reactions.

### 2. Implementation of the designed IPDO biosynthesis routes in *E. coli*

#### 2.1 Route B

As illustrated in Figure 1, route B is designed to synthesize IPDO via extending α-KIC catabolism. HPD catalyzes the transformation of α-KIC to 3-OH-IV, which is subsequently reduced to IPDO under the catalysis of CAR and PDDH, respectively. For the implementation, we began with the verification of the second step in route B, i.e. the conversion of 3-OH-IV to 3-OH-3-M-BA catalyzed by CAR. This is because that CAR has a wide scope of substrates due to its high promiscuous activity on non-native substrates (Winkler 2018). To this end, CAR from *Nocardia iowensis* was selected as the first candidate because of its wide scope of substrates (Winkler 2018). In addition, it has been reported that the 4’-phosphopantetheinyl transferases Sfp from *Bacillus subtilis* can catalyze post-translational modification of CAR by linking phosphopantetheine to the conserved serine of CAR to obtain a fully functional CAR (William Finnigan 2017). Both the *car* and *sfp* genes were synthesized and inserted into pCDFDuet, a vector that is designed for the co-expression of two target genes driven by the T7 promoter, generating thereby the plasmid pCDF-car-sfp. The latter was transformed into the *E. coli* BL21(DE3) strain to obtain a recombinant strain named IPB01 for the subsequent expression and purification of CAR. The results indicate that co-expression of CAR and Sfp is crucial for enhancing the activity of CAR (Table S3) which is in accordance with the previous study (William Finnigan 2017). For the enzyme activity assay of CAR on 3-OH-IV, benzoic acid (BA), the native substrate of CAR, was used as the positive control. As illustrated in Figure 3a, CAR showed activity on 3-OH-IV, although the activity was more 10 times lower than that on benzoic acid (0.045 *vs* 0.511μmol/min/mg).

**Figure 3.**
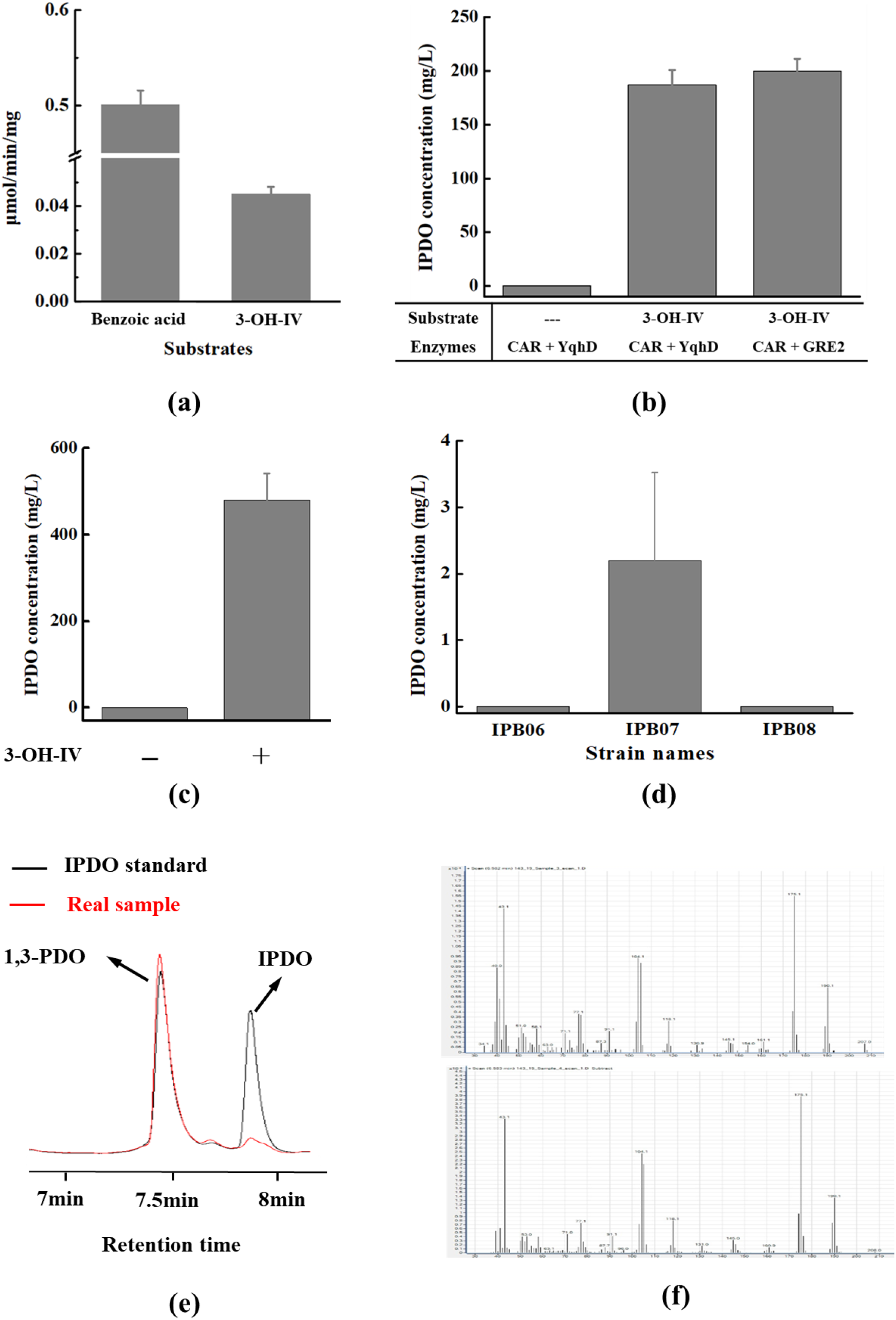
Verification of IPDO biosynthesis via route B. (a) Activity of CAR on 3-OH-IV. Benzoic acid was used as positive control. (b) Production of IPDO from 3-OH-IV by muti-enzyme cascade reaction with purified CAR and YqhD or GRE2 *in vitro*. (c) Production of IPDO from 3-OH-IV *in vivo*. Strain IPB02 was cultivated in the FM-II medium with or without addition of 50 mM 3-OH-IV into the FM-II medium. (d) IPDO production from glucose *in vivo*. (e) GC chromatograms of a calibration standard containing 100 mg/L of IPDO and a real sample from IPB07 culture. 100 mg/L 1,3-PDO was used as internal standard to monitor the derivatization efficiency with phenylboronic acid. (f) Mass spectrometry verification of IPDO production IPB07 as shown by the spectra of an IPDO standard (upper) and a real sample from IPB07 culture using glucose as sole substrate (lower). 3-OH-IV, 3-hydroxyisovalerate. 1,3-PDO, 1,3-propanediol; IPDO, isopentyldiol.

Next, the feasibility of IPDO production from 3-OH-IV was verified both *in vitro* and *in vivo*. For the *in vitro* assay, the reaction was performed in HEPES buffer (pH 7.5) containing 0 or 20 mM 3-OH-IV, 10 mM NADPH, 2 mM ATP, 10 mM Mg^2+^, 400 µg CAR, and 200 µg of either the 1,3-propanediol dehydrogenase YqhD from *E. coli* or the methylglyoxal reductase GRE2 from *Saccharomyces cerevisiae* strain S288c. The reaction was incubated at 30°C for 6 h. As shown in Figure 3b, 187.2 mg/L and 200.0 mg/L (nearly 2 mM) IPDO were formed from 3-OH-IV by the reactions containing YqhD or GRE2, respectively, while no IPDO was detected in the absence of 3-OH-IV. The higher IPDO titer of the reaction system employing GRE2 demonstrates that the exogenous GRE2 showed even a slightly higher activity on 3-OH-IV than the endogenous YqhD. Given that the amount of IPDO generated was close to the amount of ATP present in the reaction system, it is reasonable to claim that little intermediate 3-OH-3-M-BA was accumulated, and the conversion of 3-OH-IV to 3-OH-3-M-BA by YqhD or GRE2 was very efficient.

To verify the feasibility of IPDO production from 3-OH-IV *in vivo*, we ligated GRE2 into pET28a to obtain pET-gre, and co-transformed pET-gre and pCDF-car-sfp into *E. coli* BL21(DE3). The recombinant strain named IPB02 was inoculated into 5 mL LB medium and grew overnight. The overnight culture was then inoculated into 10 mL of the FM-II medium with an initial OD of 0.1 with or without the addition of 50 mM 3-OH-IV. 0.1 mM IPTG was added after OD_600_ reached 0.4 ∼ 0.6. The results showed that 480.0 mg/L IPDO were produced by IPB02 when 50 mM 3-OH-IV were supplemented in the medium, whereas no IPDO was detected in the absence of 3-OH-IV under the same conditions (Figure 3c).

Next, the synthesis of IPDO from α-KIC was verified experimentally. We used whole-cell catalysis to convert α-KIC into 3-OH-IV, as described in section 7 of the Materials and methods part. The obtained reaction mixture containing 3-OH-IV was subsequently used for the production of IPDO *in vitro* as described above. As shown in Table 2, in the negative control using *E. coli* BL21(DE3) strain no conversion of α-KIC to 3-OH-IV could be detected, while both SnHPD and AaHPD showed the α-ketoisocaproate dioxygenase activity on the conversion of α-KIC to 3-OH-IV, with AaHPD showing a slightly higher activity, as evidenced by the higher amounts of 3-OH-IV produced by IPB04 harboring AhHPD at both 6 h and 24 h compared to those produced by IPB03 harboring SnHPD (Table 2). In this regard, AhHPD was selected for further research of this pathway.

**Table 2.** Biosynthesis of IPDO from α-KIC by whole cell catalysis plus *in vitro* enzymatic reactions.

| strain | Production of 3-OH-IV from α-KIC<br>by whole cell catalysis | | | *In vitro* conversion of 3-OH-IV to<br>IPDO | |
| --- | --- | --- | --- | --- | --- |
|  | Enzymes | Time (h) | 3-OH-IV<br>(mg/L) | 3-OH-IV<br>(mg/L) | IPDO production<br>(mg/L) |
| *E. coli*<br>BL21(DE3) | No | 6 | N.D. | N.D. | N.D. |
| IPB03 | SnHPD | 6 | 914.3 | 274.3 | 232 |
|  |  | 24 | 1812.1 | 543.6 | 372 |
| IPB04 | AaHPD | 6 | 1294.7 | 388.4 | 296 |
|  |  | 24 | 2119.2 | 635.8 | 416 |

Based on the above results, we sought to explore the feasibility of producing IPDO *in vivo* from glucose. We first improved the CAR activity by optimizing the ratios of the expression levels of CAR and Sfp. The strain IPB05, which was constructed by integrating Sfp into the chromosome of *E. coli* W3110 driven by an artificial promoter J23105 (http://parts.igem.org/) and overexpressing CAR on the plasmid pTrc99a (pTrc-car), showed highest CAR activity (Table S3), so it was used for the subsequent verification of route B. Subsequently, GRE2 and AaHPD were inserted into the pTrc-car plasmid to obtain pTrc-cgh. After substituting pTrc-car by pTrc-cgh in IPB05, the resulting strain (named IPB06) was cultivated in the FM-II medium for IPDO production. Unfortunately, no IPDO was detected to be produced by IPB06. One possible reason is that the supply of precursor α-KIC is insufficient in IPB06. For this reason, we overexpressed the leucine operon (*leuA*^*fbr*^*BCD* (Tashiro et al. 2015) from *E. coli, alsS* from *B. subtilis* and *ilvCD* genes from *E. coli* on the p15AS vector to obtain the plasmid p15-leu-acd, and transformed p15-leu-acd into IPB06 to generate strain IPB07. Strain IPC08 obtained by transforming the empty p15AS vector into IPB06 was also constructed as the negative control. As the result, strain IPB07 produced 2.2 mg/L IPDO in shake flask culture, while no IPDO was produced by the control strain IPB08 (Figure 3d, 3e, 3f), demonstrating that route B is functioning *in vivo* biosynthesis of IPDO from glucose but needs to be improved in future study.

#### 2.2 Route C and D

Route C starts from α, β-DHIV and requires four exogenous reaction steps to produce IPDO. To begin with, the activity of CAR on α,β-DHIV was tested *in vitro*. As illustrated in Figure 4a, the catalytic activity of CAR on α,β-DHIV was successfully detected (0.093 μmol/min/mg), despite it was less than half of that on benzoic acid, the natural substrate of CAR. In order to perform *in vivo* validation of this route, plasmid pCOLA-yqhD-dhaB was constructed by ligating *yqhD* gene encoding an 1,3-propanediol dehydrogenase (PDDH) from *E. coli* and *dhaB* gene encoding glycerol dehydratase (GDHt) from *Klebsiella* into the pCOLADuet vector, a vector with a ColA origin for the simultaneous co-expression of two genes. Then, the plasmids pCDF-car-sfp and pCOLA-yqhD-dhaB were co-transformed into *E. coli* BL21(DE3) to generate a recombinant strain named IPC01. GC analysis showed that no IPDO production by IPC01 was detected after 48 h of cultivation. Next, through a system metabolic engineering modification of IPC01, strain IPC02 was constructed to enhance the precursor supply by increasing the flux of the α,β-DHIV biosynthesis pathway and block the branched-chain amino acid pathways in IPC01 (Figure S3). Unfortunately, strain IPC02 still didn’t produce IPDO.

**Figure 4.**
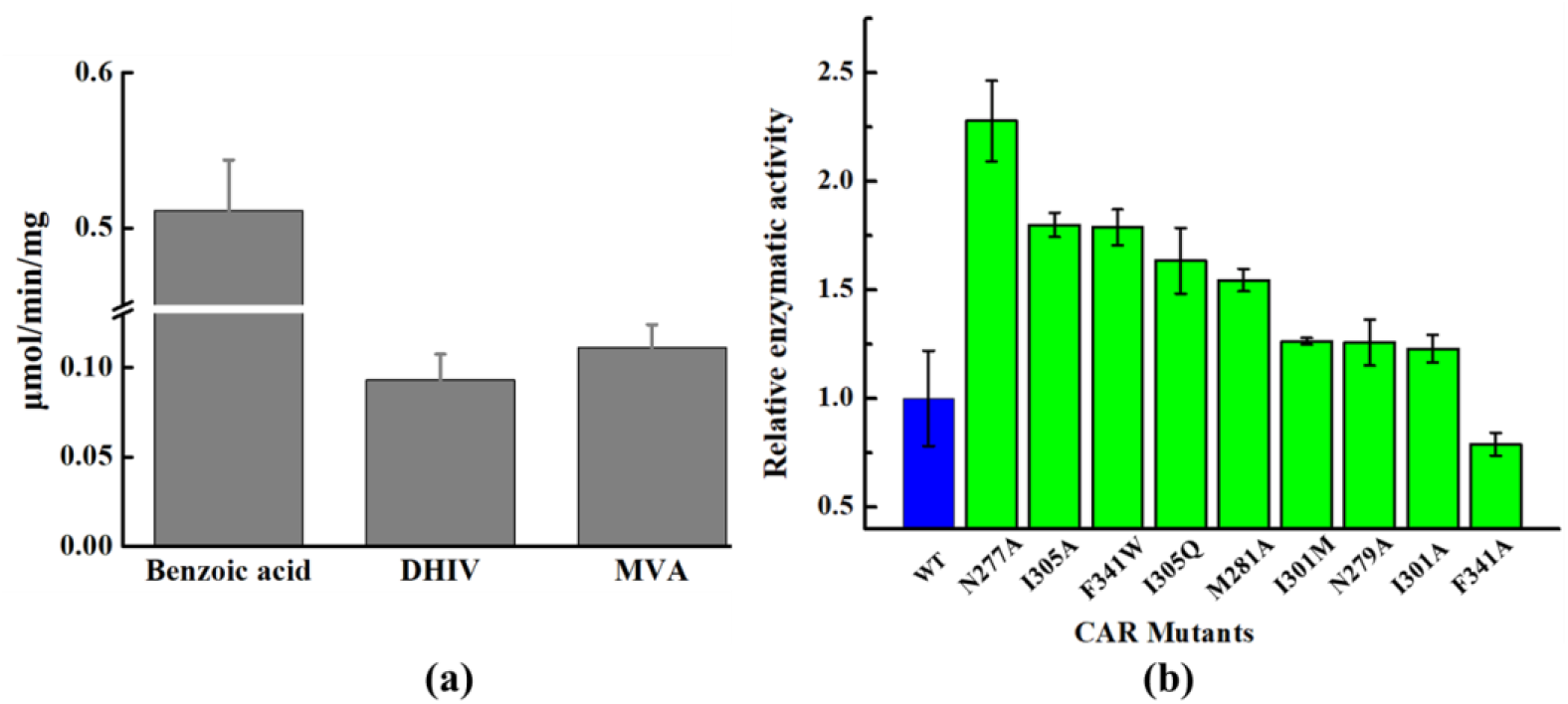
Implementation of IPDO biosynthesis via route C and D in *E. coli*. (a). Verification of the catalytic activity of CAR on α, β-DHIV and MVA *in vitro*. Benzoic acid was taken as the positive control. (b). Relative activity of CAR mutants on MVA. All CAR mutants were obtained by mutating amino acid residues near the binding pocket or on the loop region outside the binding pocket.

Then, we turned to test the feasibility of route C *in vitro*. All three enzymes CAR, DhaB and YqhD were separately purified *in vitro* and 25 mM α,β-DHIV was used as the substrate. The reactions were carried out in two steps: the first reaction catalyzed by CAR was conducted for 1 h, and then YqhD and DhaB were added into the reaction system to continue the reactions for 3 h. Unfortunately, no IPDO was detected.

Routes A to C use either a branched-chain amino acid BCAA (leucine in route A) or intermediates (α-KIC in route B and α,β-DHIV in route C) in the BCAA biosynthesis pathway and therefore extending or diverting the BCAA metabolic fluxes to the synthesis of IPDO. As an alternative route, route D is quite different in using pyruvate as the precursor for IPOD biosynthesis. It requires three reaction steps to generate mevalonate (MVA) from pyruvate and two reaction steps catalyzed successively by CAR and ADO to convert MVA to IPDO. For the implementation of route D in *E. coli*, we first verified that CAR shows activity on MVA (Figure 4a). Afterwards, the *ado* gene encoding aldehyde-deformylating oxygenase from *Prochlorococcus marinus*, as well as the *fdx* gene encoding ferredoxin from *Synechocystis* sp. 6803 and the *fnr* encoding ferredoxin (flavodoxin) NADP^+^ reductase from *E. coli* that are used to supply NADPH for the ADO catalyzed reaction, were inserted into the pCOLADuet vector to generate the plasmid pCOLA-aff. Then, the plasmids pCDF-car-sfp and pCOLA-aff were co-transformed into *E. coli* BL21(DE3) to obtain a recombinant strain named IPD01. IPD01 was cultivated in the FM-II medium by using glucose and MVA as co-substrates. According to the GC analysis results, IPD01 didn’t generate IPDO after 48 h of cultivation, even in the presence of MVA. One major reason for the failure of verification of route C and D is attributed to their poor thermodynamic driving force, especially for the CAR catalyzed reaction and two reductases catalyzed reactions in route C and the overall reactions from pyruvate to 3,5-diOH-3-M-PA in route D. In addition, the poor activities of CAR on α,β-DHIV and MVA further reduce the efficiencies of route C and D, rendering that no IPDO was formed or, at least, the generated IPDO was under the detection limit of GC analysis (less than 1 mg/L).

Next, rational engineering of CAR was carried out to improve its catalytic activity on α,β-DHIV and MVA. Computational saturation mutagenesis/virtual screening (Zhang et al. 2019) was conducted to predict CAR mutants with higher activities on MVA. Based on the crystal structure of CAR, we carried out rational protein design of CAR by either focusing on the binding pocket or on the loop near the binding pocket (Figure S4). Enzymatic activities of the predicted CAR mutants in comparison to the wild-type CAR were examined *in vitro*, as shown in Figure 4b. It’s found that most of the mutants exhibited increased activities towards MVA. Especially, the catalytic activity of the mutant N227A reached 2.28-fold of that of the wild type. Unfortunately, after these CAR mutants were introduced into route D instead of the wild-type CAR, none of the resulting recombinant *E. coli* strain showed to be able to produce IPDO *in vivo*.

## Discussion

In this study we have designed three new IPDO biosynthesis pathways in addition to the biosynthetic route (named route A in this study) we reported before (Liu et al. 2022). Among the three new routes, route B was computationally designed using retropath-RL and also successfully verified in *E. coli*, while no production of IPDO could be achieved via the two intuitively designed routes C and D. The feasibility of a synthetic pathway is controlled by many constraints, among which thermodynamics, kinetics and regulation of the enzymes involved, and concentration and physiochemical properties of metabolites (intermediates and products) and enzymes play predominant roles. In many artificial pathways, some non-natural intermediates are generated due to the promiscuity of the employed pathway enzymes, which makes it difficult to evaluate the physiochemical properties of these metabolites. The enzyme kinetic properties are always laborious and time-consuming to determine and often vary depending on the test conditions as well as on their origins, which largely limits their applications on describing the control that enzyme kinetics exerts on metabolic fluxes. As an alternative, thermodynamic analysis can be performed without using enzyme kinetic parameters and is not dependent on a particular initial steady state (Noor et al. 2015). Here, taking route A as reference route, we compared the four designed IPDO routes from the perspective of thermodynamic potential in order to better understand the implementation results. As mentioned above, the hydroxyl group can be obtained through either oxidation of an alkyl group (named oxidation pathway) or reduction of a carboxyl group to carbonyl group and then to hydroxyl group (named reduction pathway). One thing that routes B, C and D have in common and is different to route A, is that CAR is employed to catalyze the reduction of the carboxylic acid to the corresponding aldehyde using NADPH as the electron donor. However, one major drawback of these CAR-catalyzed reactions is that they operate at low rate constrained by low gibbs formation energy, especially the one in route C (Figure 1). In other words, an enzyme like CAR catalyzing a near-equilibrium reaction may “waste” almost half amount of the enzyme in catalyzing the backward reaction. In addition, the low gibbs formation energy of CAR-catalyzed reaction results in the poor thermodynamic driving force of the overall routes, as indicated by the lower MDF values of route B and C than that of route A. Therefore, to sustain the flux of the forward reaction, a large amount of enzyme and a high concentration of substrate are required which is very difficult to achieve even in the engineered IPDO strains. On top of this main thermodynamic barrier, poor enzyme kinetics of CAR, i.e. low affinity to substrate and low catalytic rate, especially *in vivo*, further restricts the metabolic flux of the pathways containing the CAR-catalyzed reactions. In contrast, the reactions that catalyze the formation of hydroxyl group through oxidation pathway are thermodynamic favorable since they are often coupled with other exergonic reactions. For instance, hydroxylation of α-KIC by oxygenase HPD in route B couples the oxidation of α-KIC with its decarboxylation, and hydroxylation of leucine by MFL in route A couples with the oxidation of α-KG to succinate. However, the high energetics of these reactions (Δ_r_G’: -100 ∼ -120 kcal/mol) comes at the expense of one carbon unit loss, which leads to the low theoretical yields of IPDO in routes A and B (Table 1). Overall, a tradeoff between the IPDO yield and titer exists in the four routes: routes carrying the oxidative pathway are highly energetic but with lower product yields, while the reductive pathway doesn’t result in carbon loss but is thermodynamically unfavorable (Figure 5). Comparing the implementation results of the four designed pathways in *E. coli*, we infer that because of the high thermodynamic driving forces of route A and B, a significant amount of metabolic flux is channeled to the IPDO synthesis pathway even at low concentrations of the precursors. In contrast, the failed implementation of route C and D is the combined result of low energetic driving force and poor kinetics of the reactions catalyzed by CAR. These results underline the significance of performing a priori thermodynamic analysis to evaluate different biosynthetic pathways that produce the same product.

**Figure 5.**
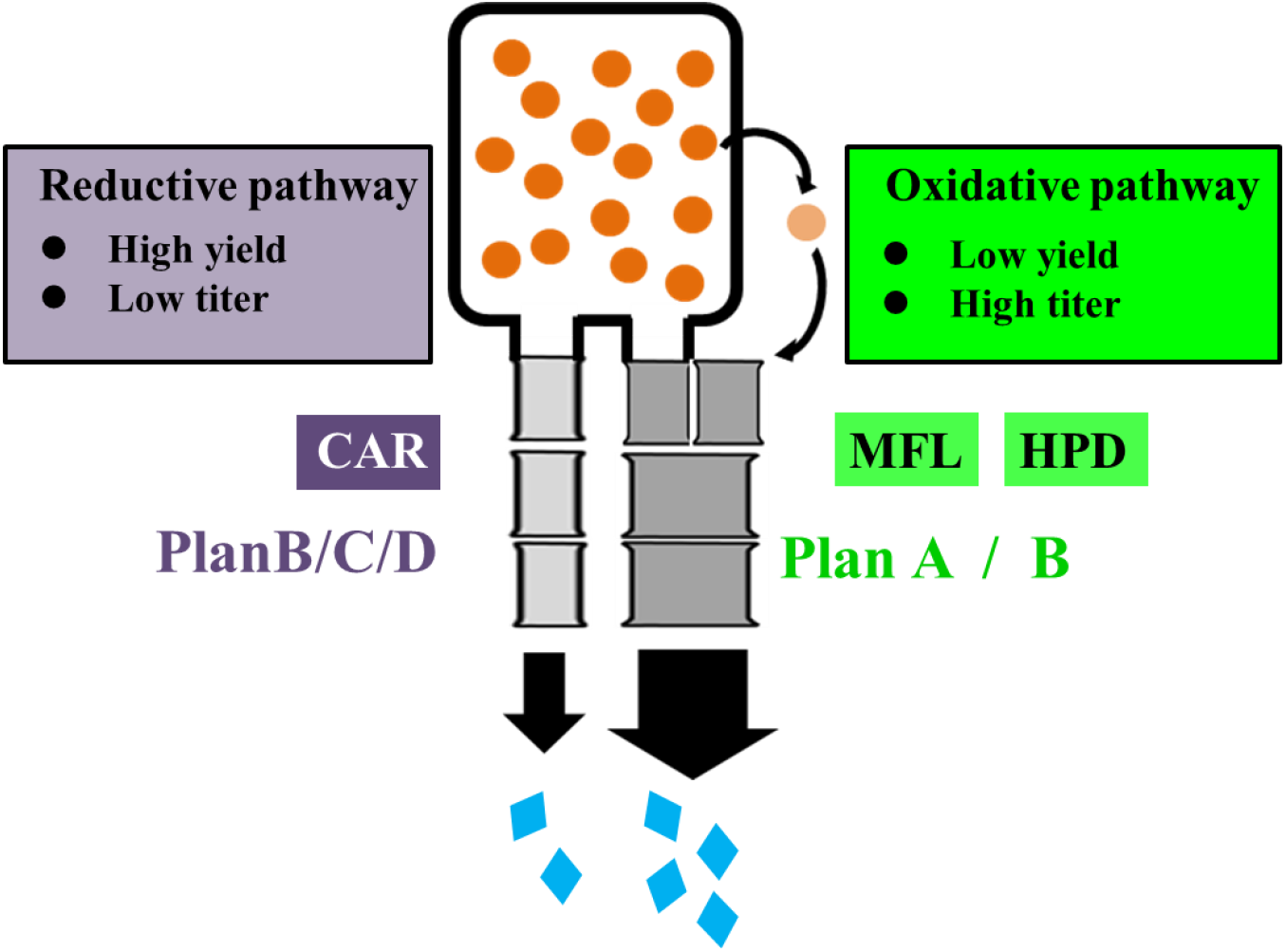
Schematic illustration of the difference in thermodynamic features between the reductive pathway and the oxidative pathway for IPDO biosynthesis. A tradeoff between the theoretical yield and the titer of IPDO exists in the four designed IPDO routes. Routes that obtain hydroxyl group through reductive pathway give higher IPDO theoretical yield but lower titer, and *vice versa*.

In addition, the efficiencies of the artificial pathways are influenced by the host strain’s intrinsic environment. These non-natural pathways are designed to synthesize target products by rewiring carbon flux from the inherent metabolisms in a competitive manner, therefore, the affinity and activity of the enzymes employed in a heterologous pathway for precursor intermediates are crucial for its efficiency. For instance, in route B, α-KIC is the precursor for IPDO biosynthesis. In wild type *E. coli* strain, the majority of the endogenous α-KIC pool is converted to leucine which is catalyzed by the branched-chain-amino-acid transaminase IlvE at high efficiency (K_m_: 0.08 mM, K_cat_: 24.7 s^-1^) (Yu et al. 2014). In contrast, the catalytic ability of HPD on α-KIC is much lower (K_m_: 0.32 mM, K_cat_: 6.5 min^-1^) (Sabourin and Bieber 1982) than that of IlvE. Given this, it is less efficient to use HPD to divert the endogenous α-KIC from leucine synthesis pathway into IPDO synthesis pathway (Table 1). This is also the case for route C, in which CAR competes with endogenous dihydroxy-acid dehydratase IlvD for binding the substrate α,β-DHIV for IPDO synthesis (Table 1). For further improve the efficiency of route B in future work, on the one hand, the activity of HPD on α-KIC should be significantly improved through either rational protein engineering or screening HPDs with higher activities from different species. On the other hand, a dynamic regulation system can be employed to dynamically control the expression abundance of IlvE. Recently Hou *et al*. constructed a bifunctional molecular switch system by using growth phase-dependent promoters and degrons(Hou et al. 2020). Within their system, dynamic regulation of the expression of IlvE at different growth phases can be fulfilled: A high expression level is kept at the exponential growth phase to ensure good cell growth, but turned down to a relatively low level when cells reach the stationary phase to avoid shunting metabolic flux towards valine and leucine synthesis. In addition, a series of promoters with different strengths and induction phases have been characterized in their toolbox, which can be adopted in our future study for fine-tuning the IlvE expression to maximize IPDO production.

## Conclusion

In this study we provided a framework to evaluate the potential of four different IPDO biosynthesis routes prior to experimental implementation. Several parallel criteria, including the theoretical yield, thermodynamics, energy requirement and biochemical properties of intermediates and enzymes, were used to explore the potential efficiencies of different pathways prior to implementation. By integration of *in silico* analysis results, route A ranked the highest among the four pathways: it shows a higher thermodynamic driving force, requires least ATP and reducing power and has no toxic intermediates or enzymes with unfavorable properties. To demonstrate the applicability of this evaluation framework, we implemented all four artificial IPDO pathways using *E. coli* as the host strain and compared their performance with the evaluation results. The results revealed that route A, which ranks the highest according to the *in silico* analysis, shows the best performance on the IPDO production *in vivo*. The consistency of the theoretical analysis (dry test) and wet test proves that such an integration of tools is very useful in developing and analyzing the novel artificial pathway designs.

## Acknowledgements

We thank Hua An Tang Company (China) for providing financial support for this study. YL is supported by a PhD scholarship from the Chinese Scholarship Council (CSC), which is gratefully acknowledged.

## Author contributions

AZ, YL, LC and CM conceived the concept of the work. YL, LC, CM and QY designed and performed the experimental studies. YL, LC and WW did the analytic work. YL, LC, WW, QY, HM, CZ, CM and AZ involved in data analysis and discussion and in writing the manuscript. AZ provided the financial support and supervised the work.

