## Supplemental material for "Design, evaluation and implementation of synthetic isopentyldiol pathways in *Escherichia coli*"

*”*

Yongfei Liu^1#^, Lin Chen^1#^, Pi Liu^2^, Qianqian Yuan^2^, Chengwei Ma^1^, Wei Wang^1^, Chijian Zhang^1，3^, Hongwu Ma^2^ and An-Ping Zeng^1,4,*^

^1^Hamburg University of Technology, Institute of Bioprocess and Biosystems Engineering, Denickestr. 15, Hamburg 21073, Germany

^2^Tianjin Institute of Industrial Biotechnology, Chinese Academy of Sciences, Tianjin 300308, China

^3^Hua An Tang Biotech Group Co., Ltd, Guangzhou, China

^4^Present address: Center of Synthetic Biology and Integrated Bioengineering, School of Engineering, Westlake University, Hangzhou, Zhejiang, 310024, China, 310024

^#^ These authors contribute equally to this article.

Figure S1. Two IPDO biosynthesis routes designed using the software RetropathRL. α-KIC, α-ketoisocaproate; 3-OH-IV, 3-hydroxyisovalerate; 3-OH-3-M-BA, 3-hydroxy-3-methylbutanal; IPDO, isopentyldiol; E-3-OH-IV: ethyl 3-hydroxy-isovalerate.

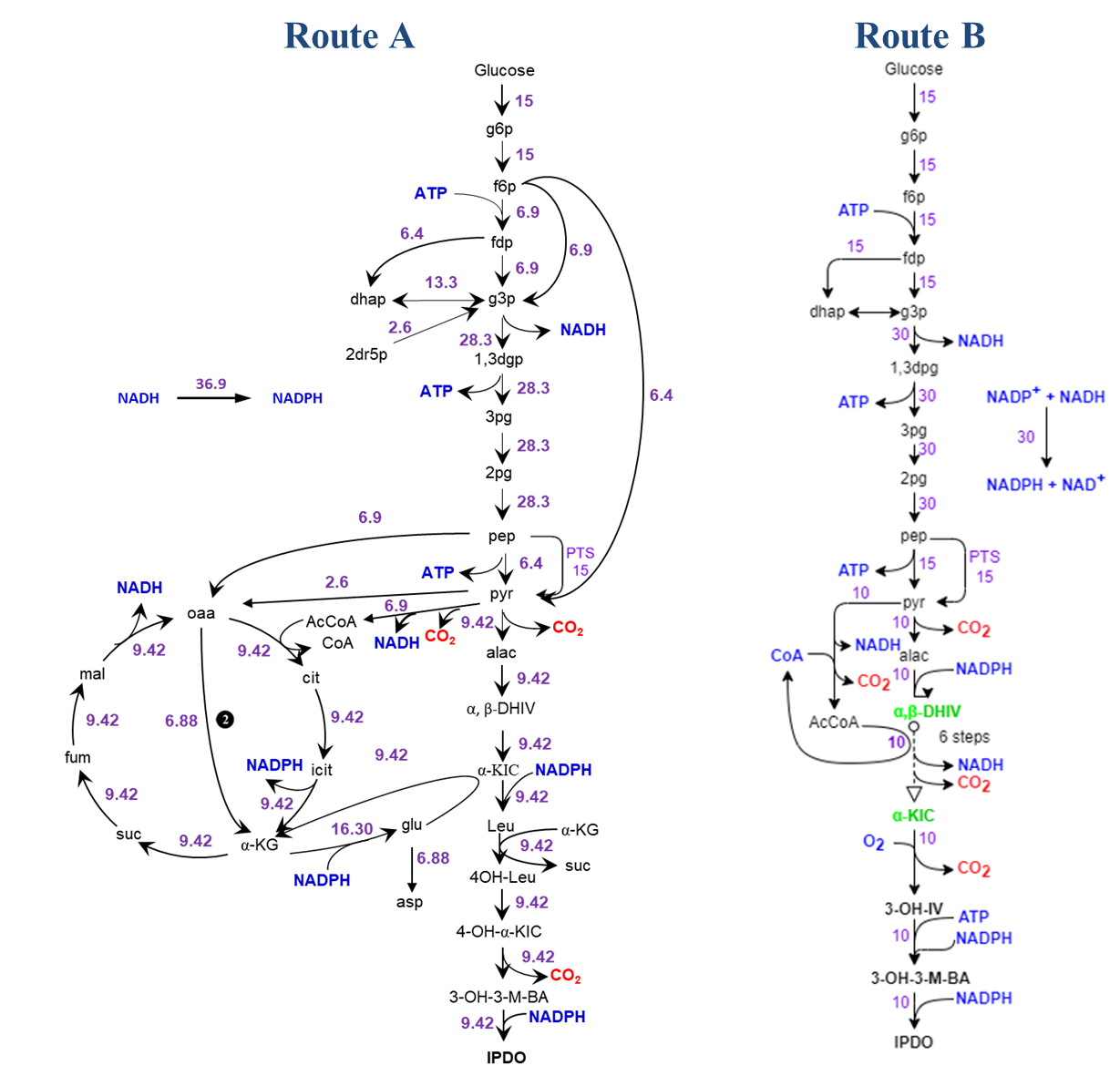

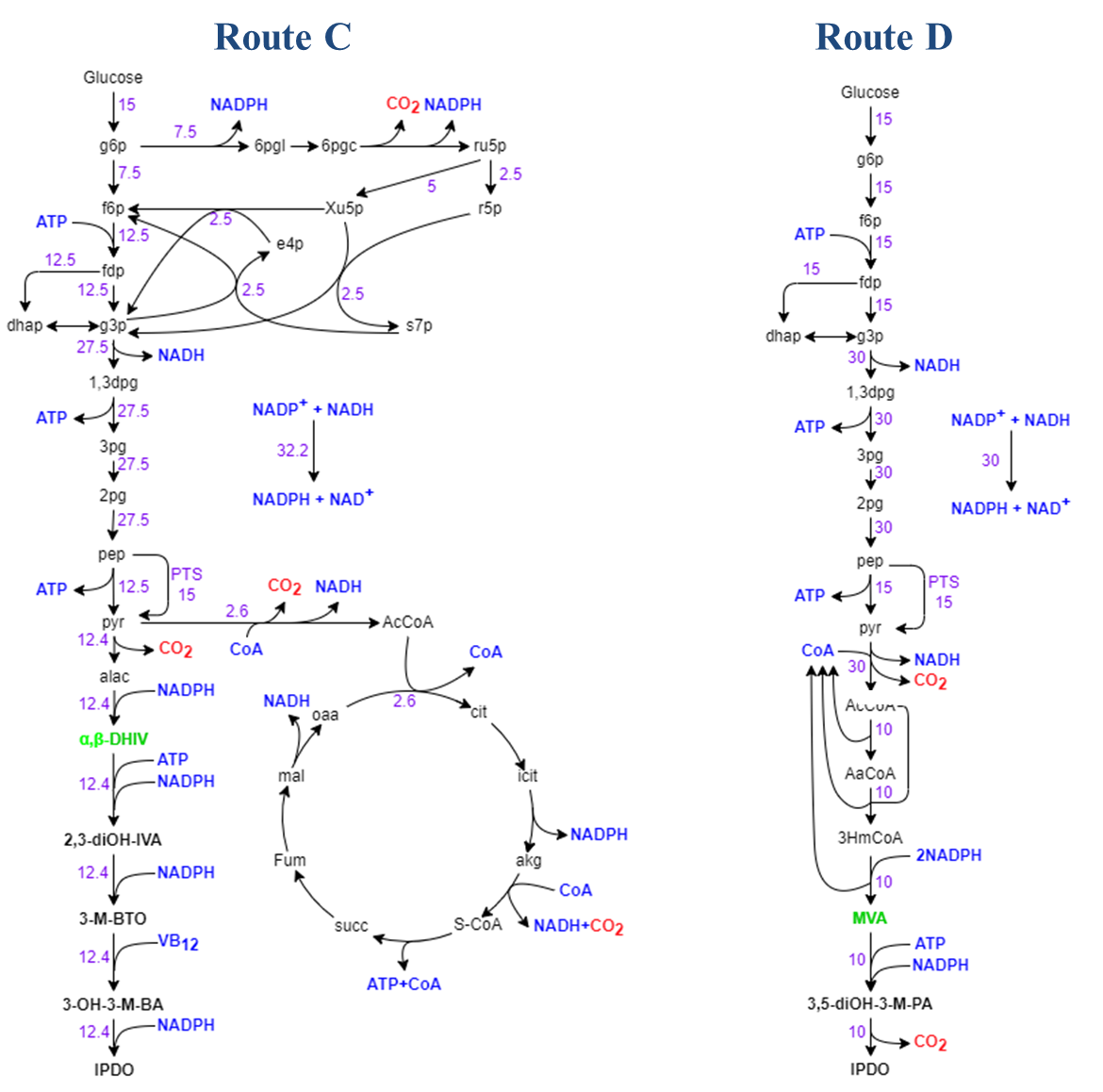

Figure S2. Optimal metabolic flux distribution for IPDO production by four different artificial routes in *E. coli* iY75_1357 odel. Abbreviations: g6p, glucose-6-phosphate; f6p, fructose-6-phosphate; fdp, fructose-1,6-bisphosphate; g3p, glyceraldehyde-3-phosphate; 1,3-dgp, 1,3-bisphosphoglycerate; 3pg, glycerate-3-phosphate; 2pg, glycerate-2-phosphate; pep, phosphoenolpyruvate; pyr, Pyruvate; AcCoA, acetyl coenzyme A; oaa, oxalosuccinate; cit, citric acid; cis-aco, cis-aconitase; icit, isocitrate; akg, α-ketoglutarate; S-CoA, succinyl-CoA; succ, succinate; fum, fumarate; mal, malate; 6pgi, 6-phosphate-glucose lactone; ru5P, ribulose-5-phosphate; Xu5P, xylulose-5-phosphate; r5p, ribose-5-phosphate; s7p, sedoheptulose-7-phosphate; e4p, erythrose-4-phosphate; ATP, adenosine triphosphate; ADP, adenosine diphosphate; NADPH, nicotinamide adenine dinucleotide phosphate; NADH, Nicotinamide adenine dinucleotide; 4OH-Leu, 4-hydroxyleucine; 4-OH-α-KIC, 4-hydroxy-4-methyl-α-ketovaleric acid; 3-OH-3-M-BA, 3-hydroxy-3-methylbutanal; α-KIC, α-ketoisocaproate; 3-OH-IV, 3-hydroxyisovalerate; α, β-DHIV, α, β-dihydroxyisovalerate; 2,3-diOH-IVA, 2,3-hydroxy-3-methylbutanal; 3-M-BTO, 3-methyl-1,2,3-butanetriol; MVA, mevalonate; 3,5-diOH-3-M-PA, 3,5-dihydroxy-3-methylpentanal; IPDO, isopentyldiol.

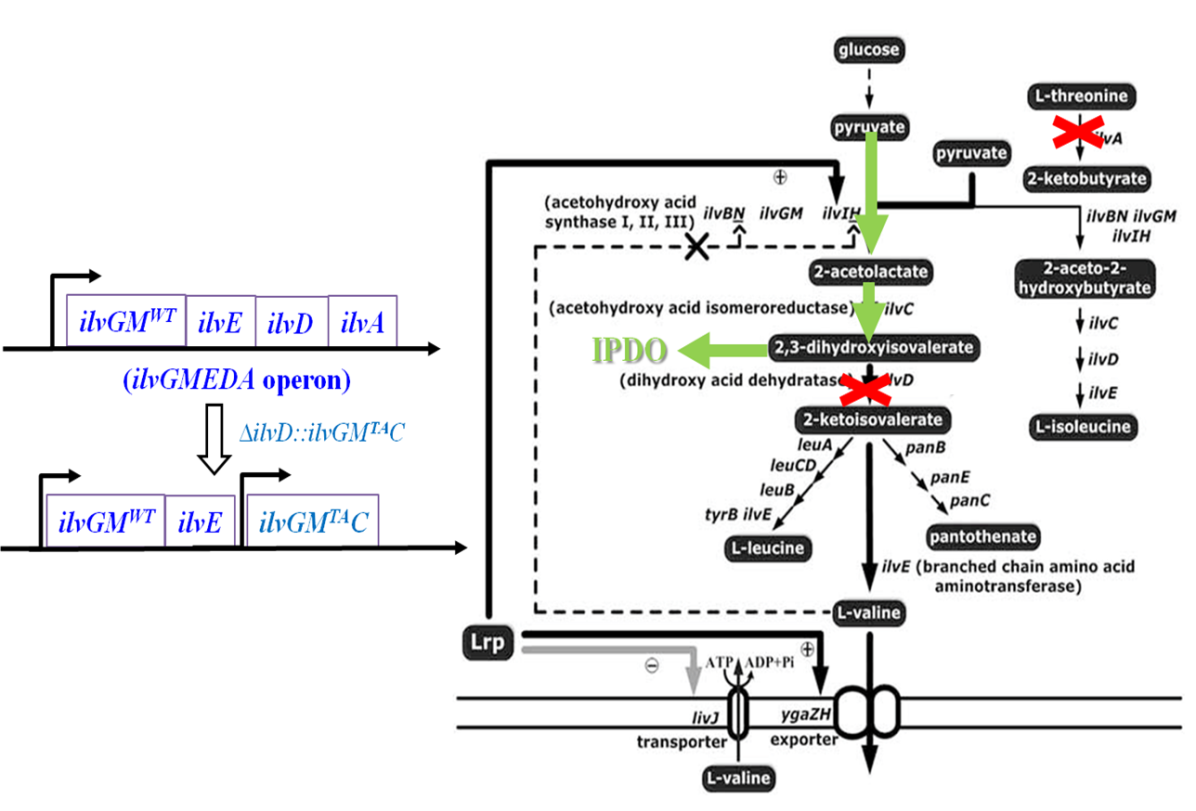

Figure S3. System metabolic engineering on *E. coli* BL21(DE3) to enhance the accumulation of α, β-DHIV. Gene *ilvD* and gene *ilvA* were replaced by *ilvGM*^TA^ and *ilvC* via genomic editing. The gene *ilvGM*^TA^ was obtained by repairing the frame-shift mutation existing in the wild type of *ilvGM* (*ilvGM*^WT^). Thus, enhancement of the gene expression for *ilvGM*^TA^ and *ilvC* increased the metabolic flux from pyruvate to α, β-DHIV and thus provided more precursors for the IPDO biosynthesis.

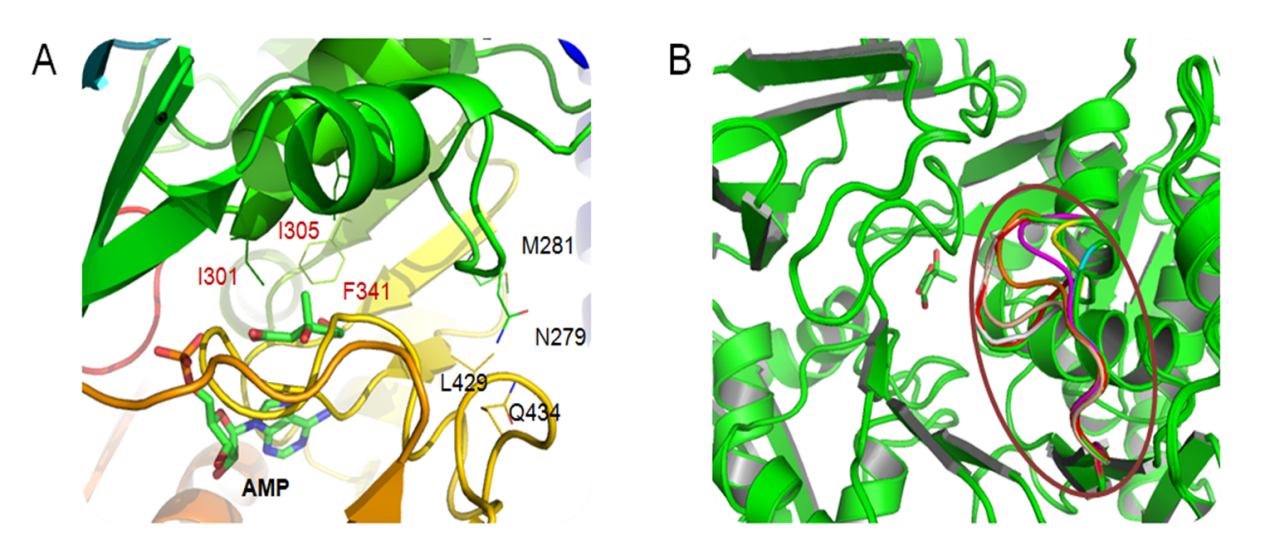

Figure S4. Rational protein design of CAR. (A) Focused on the binding pocket. The crystal structure of CAR was retrieved from the Protein Data Bank (PDB, entry No: 5msd). For the mutation sites located in the binding pocket, alanine-scanning mutagenesis was first suggested for sites I301, F341 and I305. Then, computational saturation mutagenesis/virtual screening was conducted and mutations I301M F341W I305Q were predicted (A). (B) Focused on the loop outside the binding pocket. For the loop outside of the binding pocket, alanine-scanning mutagenesis is suggested for M281, L429, N279 Q434, Q277, Q283, N285, Q287 and R288. Then loop engineering was suggested for the fragment NSMLQGNSQRVG (indicated by the red circle) in which the first six residues were replaced by different length of poly(Gly).

Table S1. Strains and plasmids used in this work

| Strains / plasmids | Description | Source or reference |
| --- | --- | --- |
| Plasmids |  |  |
| p15AS |  | Fang et al*^1^* |
| pCOLADuet |  |  |
| pCDFDuet |  |  |
| pCas-Red | Applied for genomic editing | Dongdong Zhao*^2^* |
| pZA | Deleting luciferase gene from pZA31-Luc | Stored in our lab*^3^* |
| pZA-aky | pZA carrying *aad*, *kdc* and *yqhD* genes | This work |
| pZA-akym | pZA carrying *aad*, *kdc,* *yqhD* and *mfl* genes | This work |
| pET-mfl | pET28a carrying *mfl* gene | This work |
| pET-car | pET28a carrying *car* gene | This work |
| pCDF-car-sfp | pCDFDuet carrying *sfp* and *car* gene | This work |
| pET-gre2 | pET28a carrying *gre2* gene | This work |
| pET-snhpd | pET22b carrying *snhpd* gene from *Streptomyces noursei* | This work |
| pET-aahpd | pET22b carrying *aahpd* gene from *Allokutzneria albata* | This work |
| pTrc-cgh | pTrc99a carrying *aahpd*, *car* and *gre2* gene | This work |
| p15-leu-acd | p15AS carrying *alsS*, *ilvCD* and *leuA^fbr^BCD* genes | This work |
| pCOLA-yqhD-dhaB | pCOLADuet carrying *yqhD* and *dhaB* genes | This work |
| pET-yqhd | pET28a carrying *yqhD* gene | This work |
| pET-dhaB | pET28a carrying *dhaB* gene | This work |
| pCOLA-aff | pCOLADuet carrying *ado*, *fdx* and *fnr* genes | This work |
| Strains |  |  |
| *E. coli* Top10 |  | New England Biolabs |
| *E. coli* BL21 (DE3) |  | New England Biolabs |
| *E. coli* DY330 | W3110 ΔlacU169 gal490 λCI857 Δ(*cro-bioA*) | Yu et al*^4^* |
| IPA00 | *E. coli* BL21 (DE3) carrying pZA | This work |
| IPA01 | *E. coli* BL21 (DE3) carrying pZA-aky | This work |
| IPA02 | *E. coli* BL21 (DE3) carrying pZA-maky | This work |
| IPA03 | *E. coli* BL21 (DE3) carrying pZA-aky and pET-mfl | This work |
| IPB00 | *E. coli* BL21 (DE3) carrying pET-car | This work |
| IPB01 | *E. coli* BL21 (DE3) carrying pCDF-car-sfp | This work |
| IPB02 | *E. coli* BL21 (DE3) carrying pET-gre and pCDF-car-sfp | This work |
| IPB03 | *E. coli* BL21 (DE3) carrying pET-snhpd | This work |
| IPB04 | *E. coli* BL21 (DE3) carrying pET-aahpd | This work |
| IPB05 | Chromosome integration of Sfp driven by J23105 promoter | This work |
| IPB06 | IPB05 carrying pTrc-cgh | This work |
| IPB07 | IPB06 carrying p15-leu-acd | This work |
| IPB08 | IPB06 carrying p15AS | This work |
| IPB09 | Chromosome integration of Sfp driven by T7 promoter, carrying pET-car | This work |
| IPC01 | *E. coli* BL21 (DE3) carrying pCDF-car-sfp and pCOLA-yqhD-dhaB | This work |
| IPC02 | System engineering of IPC01 to enhance the α, β-DHIV supply | This work |
| IPD01 | *E. coli* BL21 (DE3) carrying pCDF-car-sfp and pCOLA-aff | This work |

Table S2. Primers used in this work

| Names | Sequences |
| --- | --- |
| pZA-vec-LR | ctttctcctctttaatgaattcggtc |
| pZA-vec-LF | tgactctagaggcatcaaataaaac |
| pZA-aad-LF | attcattaaagaggagaaagatgaaaatcagccgccgca |
| pZA-aad-LR | atctccttcttaaagttgttactttttgaagcgatccatgct |
| pZA-kdc-LF | caactttaagaaggagatatacatatgtataccg |
| pZA-kdc-LR | tgtatatctccttcttaaagttttatttgttctgctcggcaaac |
| pZA-yqhD-LF | aactttaagaaggagatatacatatgaacaactttaatctgcacaccc |
| pZA-yqhD-LR | tttgatgcctctagagtcattagcgggcggcttcgtata |
| pZA-akym-mfl-LF | gtataatgctagcctgcagaaggagatatagtgccgcgcggcagccat |
| pZA-akym-mfl-LR | ttacaacttggcagtgaacgtc |
| pZA-akym-vec-LF | ttcactgccaagttgtaacgctacgctcggtcgttcg |
| pZA-akym-vec-LR | tgcaggctagcattatacctaggactgagctagctgtcaaagtcagtgagcgaggaagcg |
| pET-mfl-LF | gtgccgcgcggcagccatatggaggttagcgagaagct |
| pET-mfl-LR | gagctcgaattcggatccttacagcttggcggtgaacg |
| pET-vec-LF | ggatccgaattcgagctccgtcga |
| pET-vec-LR | atggctgccgcgcggcacc |
| pCDF-sfp-LF | atgggcaagatttacggtatctat |
| pCDF-sfp-LR | ttacagcagttcttcgtagctaac |
| pCDF-vec1-LF | tagctacgaagaactgctgtaaggatccgaattcgagctcgg |
| pCDF-vec1-LR | atgtatatctccttcttatacttaacta |
| pCDF-vec2-LF | gctgcaactgctgtaacctcgagtctggtaaagaaaccgctgc |
| pCDF-vec2-LR | tagataccgtaaatcttgcccatggtatatctccttattaaagttaaacaaaatt |
| pCDF-car-LF | ttaagtataagaaggagatatacatatggcggtggatagcccggat |
| pCDF-car-LR | gttacagcagttgcagcagttcc |
| pET-gre2-LF | catcaccatatgtcagttttcgtttcaggtgctaacg |
| pET-gre2-LR | ggccgcaagcttatattctgccctcaaattttaaaatttgggagg |
| pET-vec-LF | gcagaatataagcttgcggccgcactcgagca |
| pET-vec-LR | aaaactgacatatggtgatgatgatgatgatggcccatggtatatctccttcttaaa |
| pET-snhpd-LF | ctctgccatatggcagacaccacgatgcac |
| pET-snhpd-LR | attagactcgaggaggttgccgcgccggtcct |
| pET-aahpd-LF | atgatcctcgaggagattgcctcggcgctcct |
| pET-aahpd-LR | gatatacatatgaccgagactctcgaccagg |
| pTrc-car-LF | cagcctgcgtctgctgccgcgtt |
| pTrc-car-LR | cgaattccagcttgtcgacggagctcgaattcggatccttacagcagttgcagcagttc |
| pTrc-gre-LF | ttcgagctccgtcgacaagctggaattcgagctcagaaggag |
| pTrc-gre-LR | gattaattgtcaacaggtaccttatattctgccctcaaattttaaaatttgggagg |
| pTrc-hpd-LF | tcccggggatccgaaataattttgtttaactttaagaaggagatatacatatg |
| pTrc-hpd-LR | atccgccaaaacagccaagctttagagattgcctcggcgc |
| p15-leu-LF | gcagaaggagatataatgagccagcaagtcattatttt |
| p15-leu-LR | ttaattcataaacgcaggttgttt |
| p15-vec1-LF | tgcgtttatgaattaactgcaggaattcggatcccc |
| p15-vec1-LR | ctcgagactagtcatatggtaccgtt |
| p15-ilv-LF | agtctcgagttaaccccccagtttcgatt |
| p15-als-LR | gcagaaggagatatattgacaaaagcaacaaaagaacaaaaatccctt |
| pCOLA-yqhD-LF | aggagatataccatgggcaacaactttaatctgcacaccc |
| pCOLA-yqhD-LR | gtggtggtggtggtgctcgaggtggtggtggtggtggtggc |
| pCOLA-vec1-LF | ctcgagcaccaccaccaccaccactaaggatccgaattcgagctcgg |
| pCOLA-vec1-LR | tatgtatatctccttcttatactt |
| pCOLA-dhaB-LF | aaggagatatacatatgaaaagatcaaaacgatttgcag |
| pCOLA-dhaB-LR | ctttaccagactcgagttagcttcctttacgcagc |
| pCOLA-vec2-LF | tcgagtctggtaaagaaac |
| pCOLA-vec2-LR | catggtatatctccttattaaagtt |
| pCOLA-ado-LF | ggagatataccatgggcagcagcatgggcccgaccctggaaat |
| pCOLA-ado-LR | cgcgccgagctcgaattcggatccttagctaaccagcgccgccgccgccatac |
| pCOLA-fnr-LF | gaaggagatatacatatggctgattgggtaacagg |
| pCOLA-fnr-LR | ttaccagtaatgctccgctgtc |
| pCOLA-fdx-LF | acagcggagcattactggtaaagaaataattttgtttaactttaagaagga  gatatacatatggcgagctacaccgtgaaa |
| pCOLA-fdx-LR | tccaattgagatctgccatttagtacaggtcttcttctttgt |

Table S3. Comparison of the activities of CAR overexpressed from different hosts.

| Host | | Genome editing | plasmid(s) | nmol/mg/min |
| --- | --- | --- | --- | --- |
| *E. coli* BL21 |  | | pET-car | 100±38 |
| *E. coli* BL21 |  | | pCDFDeut-car-sfp | 329±7 |
| *E. coli BL21* | Chromosome integration of Sfp driven by T7 promoter | | pET-car | 470±44 |
| *E. coli* BL21 | Chromosome integration of Sfp driven by T7 promoter | | pTrc-car | 1251±184 |
| *E. coli* W3110 | Chromosome integration of Chromosome integration of Sfp driven by J23101 promoter | | pTrc-car | 1200±86 |
| *E. coli* W3110 (IPB05) | Chromosome integration of Sfp driven by J23105 promoter | | pTrc-car | 1384±190 |
